# Inductive Inference of Gene Regulatory Network Using Supervised and Semi-supervised Graph Neural Networks

**DOI:** 10.1101/2020.09.27.315382

**Authors:** Juexin Wang, Anjun Ma, Qin Ma, Dong Xu, Trupti Joshi

## Abstract

Discovering gene regulatory relationships and reconstructing gene regulatory networks (GRN) based on gene expression data is a classical, long-standing computational challenge in bioinformatics. Computationally inferring a possible regulatory relationship between two genes can be formulated as a link prediction problem between two nodes in a graph. Graph neural network (GNN) provides an opportunity to construct GRN by integrating topological neighbor propagation through the whole gene network. We propose an end-to-end gene regulatory graph neural network (GRGNN) approach to reconstruct GRNs from scratch utilizing the gene expression data, in both a supervised and a semi-supervised framework. To get better inductive generalization capability, GRN inference is formulated as a graph classification problem, to distinguish whether a subgraph centered at two nodes contains the link between the two nodes. A linked pair between a transcription factor (TF) and a target gene, and their neighbors are labeled as a positive subgraph, while an unlinked TF and target gene pair and their neighbors are labeled as a negative subgraph. A GNN model is constructed with node features from both explicit gene expression and graph embedding. We demonstrate a noisy starting graph structure built from partial information, such as Pearson’s correlation coefficient and mutual information can help guide the GRN inference through an appropriate ensemble technique. Furthermore, a semi-supervised scheme is implemented to increase the quality of the classifier. When compared with established methods, GRGNN achieved state-of-the-art performance on the DREAM5 GRN inference benchmarks. GRGNN is publicly available at https://github.com/juexinwang/GRGNN.

**Highlights:**

1. We present a novel formulation of graph classification in inferring gene regulatory relationships from gene expression and graph embedding.
2. Our method leverages a powerful framework, gene regulatory graph neural network (GRGNN), which is flexible and powerful to ensemble statistical powers from a number of heuristic skeletons.
3. Our results show GRGRNN outperforms previous supervised and unsupervised methods inductively on benchmarks.
4. GRGNN can be interpreted and explained following the biological network motif hypothesis in gene regulatory networks.

## 1. Introduction

Gene regulatory networks (GRNs) represent the causal regulatory relationships between transcription factors (TFs) and their gene targets (Marbach *et al*., 2012). Integrating sufficient regulatory information as a graph, GRNs are essential tools for elucidating gene functions, interpreting biological processes, and prioritizing candidate genes for molecular regulators and biomarkers in complex diseases and traits analyses (Marbach *et al*., 2012). While high-throughput sequencing and other post-genomics technologies enable statistical and machine learning methods to reconstruct GRN, inferring gene regulatory relationships between a set of TFs and a set of potential gene targets through gene expression data is still far from being resolved in bioinformatics (Mochida *et al*., 2018).

With decades of efforts of inferring gene regulatory relationships from gene expression data, many machine learning and statistical methods have been proposed for reconstructing GRN (Mochida *et al*., 2018). Unsupervised methods dominate GRN inference. These methods include 1) regression-based methods, in which TFs are selected by target gene through sparse linear-regression, such as TIGRESS (Haury *et al*., 2012); 2) information-based methods, such as ranking edges based on variants of mutual information, e.g. CLR (Faith *et al*., 2007); 3) correlation-based methods, such as the absolute value of Pearson’s correlation coefficient and Spearman’s correlation coefficient; and 4) Bayesian networks by optimizing posterior probabilities using different heuristic searches (Aliferis *et al*., 2010). Among all unsupervised methods, GENIE3 (Huynh-Thu *et al*., 2010) is a well-established and widely accepted method based on ensemble random forest regression of gene expression levels between TF and targets. In the DREAM5 challenge on gene network inference (Marbach *et al*., 2012), GENIE3 obtained the best performance among all the methods at that time.

In recent years, due to the identification of a large number of TFs and their targets, supervised approaches have been developed to train classifiers to infer regulatory interactions. Many studies have demonstrated that carefully trained supervised models outperform unsupervised methods (Maetschke *et al*., 2013). These supervised methods decompose the gene regulatory network inference problem into a large number of subproblems to estimate local models for characterizing the genes regulated by each TF (Maetschke *et al*., 2013). Bleakley et al. firstly reconstructed biological networks using local models in SVM (Bleakley *et al*., 2007). Other SVM-based methods include SIRENE (Mordelet and Vert, 2008) and CompareSVM (Gillani *et al*., 2014). Cerulo et al. used a probability estimation approach to learn GRN from only positive and unlabeled data (Cerulo *et al*., 2010).

With recent advancements in deep learning, there is already some work to predict gene regulatory relationships through the deep learning framework. Daoudi and Meshoul trained a deep neural network on known TF and target pairs in each of DREAM4 multifactorial data (Daoudi and Meshoul, 2019). MacLean trained a shallow convolutional neural network with known Arabidopsis TFs and target pairs with microarray gene expression as the features (MacLean, 2019). Turki et al. used unsupervised methods to train supervised models to guide SVM and deep neural networks to infer GRNs through link prediction (Turki *et al*., 2016).

However, these existing supervised GRN inferring methods show limited usage in practical biological applications. Because of heterozygous data sources, these supervised methods usually have limited generalizable capabilities in complex biological mechanisms. Most supervised models are formulated as the matrix complementation problem. All the results are based on training and testing on single data source splitting into training/validation/testing datasets or in cross validation. For a practical GRN inferring problem, there is usually no known relationship ready for training, which makes it unfeasible to predict gene regulatory relationships inductively in practice.

Moreover, gene regulatory activities always act as a whole system with a set of genes to perform a biological function (Long *et al*., 2008). Network motif (Alon, 2007) is a widely accepted biological hypothesis, that a small set of recurring regulation patterns can serve as basic building blocks of GRN. The same network motifs have been found in diverse organisms from bacteria to humans. However, these existing supervised GRN inferring methods usually only take the two endpoints of the regulatory interactions as the input, and then treat these known TF/target gene interactions independently in the training processes, and hence neglect the global relationships among these interactions. One of the related work to inductively infer GRN is by Patel and Wang (Patel and Wang, 2015). Based on SVM, they only trained and tested 4 TFs with the largest degrees with inductive and transductive inferences.

Instead of learning only two ends of the relationships, graph models are capable of modeling complex relationships between TF/gene pairs and their neighbors. Graph neural networks (GNN) as a generalization of neural networks are designed to handle graphs and graph-related problems as node classification, link prediction, and graph classification (Hamilton *et al*., 2017). Generally, GNNs consist of an iterative process to propagate the node information. After *h* iterations of the aggregation, each node in the graph can be presented by a feature vector aggregating from its *h*-hop neighbors. The entire graph can be represented by pooling on all feature vectors of all nodes in the graph.

In the context of graph analysis, link prediction is one of the major research areas of GNN (Veličković *et al*., 2017). Predicting links through an auto-encoder or variational auto-encoder achieved great success transductively (Kipf and Welling, 2016a) (Kipf and Welling, 2016b). Zhang and Chen firstly extracted local enclosed subgraphs around links to train a fully connected neural network in Weiseiler-Lehman Neural Machine (Zhang and Chen, 2017), and then SEAL (Zhang and Chen, 2018) was proposed to use a GNN to replace the fully connected neural network.

Inspired by SIRENE (Mordelet and Vert, 2008), we extended SEAL by formulating the GRN inference problem as a graph classification problem and propose an end-to-end framework gene regulatory graph neural network (GRGNN) to infer GRN. The basic hypothesis is that the features of two nodes and their neighbors (local structure) can decide whether they form a TF and target gene pair, which is consistent with the network motif hypothesis. The local structure as a graph consists of gene pairs and neighbors, which can be distinguished through a classifier. For an unknown condition or species, the biggest advantage in this formulation is inductive learning, i.e., GRNs can be constructed with the same input as the unsupervised methods without using new labels in the new condition or species.

The major innovation in this paper is introducing heuristic starting skeletons for inductive learning. The initial graph of genes is built from one of several noisy skeletons based on different heuristics on gene expression data. Then, the subgraphs centered at known TF/gene pairs are extracted. A linked pair between a TF and its target gene, and their neighbors are labeled as a positive subgraph, while an unlinked TF/target gene pair and their neighbors are labeled as a negative subgraph. GNN classifiers are trained through these subgraphs, and then ensemble together to predict links as graph labels in GRN. A semi-supervised framework is also adopted to handle the unlabeled data.

To the best of our knowledge, this is the first work to infer gene regulatory network through graph neural networks. Our contributions in this paper are (1) introduction of a supervised/semi-supervised graph classification framework for gene regulatory network inference, (2) using noisy starting skeleton to guide link prediction in the graph, and (3) efforts of inductive inference GRN across different species and conditions.

## 2. GRGNN Framework

Inferencing regulatory relationships in GRN can be defined as follows: given a set of TFs *T*, a set of target genes *G*, and gene expression data *EX*_*i, j*_, *i* ∈{*T,G*}, *j* ∈[1, *n*] for all *T* and *G* with *n* arrays, infer the adjacency matrix *A*_*T, T* +*G*_ for all *T*. GRN is defined as a bipartite graph < *T, G, E* >, where both *T* and G are vertices in the graph. *E* is the set of links in GRN. Link *E* only exists between (*T,T*) and (*T,G*), any (*G,G*) ∉ *E*. In the adjacency matrix *A, A*_*i, j*_ = 1 if (*i, j*) ∈ *E* and *A*_*i, j*_ = 0 otherwise. With the abundance of information of vertices, *X*_*i*_ is the node information corresponding to a single node *i. d*(*x, y*) is the shortest distance between node *x* and *y*. Node *x* is node *y*’s *h*-hop neighbor when *d*(*x, y*) = *h*. As this study aims to predict link existence, *E* is always treated as an undirected edge in our formulation.

Framework GRGNN is proposed to solve this problem. Figure 1 shows the scheme of GRGNN. The whole processes of GRGNN consist of the following four steps: 1) construct noisy starting skeletons; 2) extract enclosed subgraphs; 3) add node labels and features; 4) build ensemble GNN classifiers. Finally, a semi-supervised learning framework is proposed to deal with the unlabeled links in GRN.

**Figure 1.**
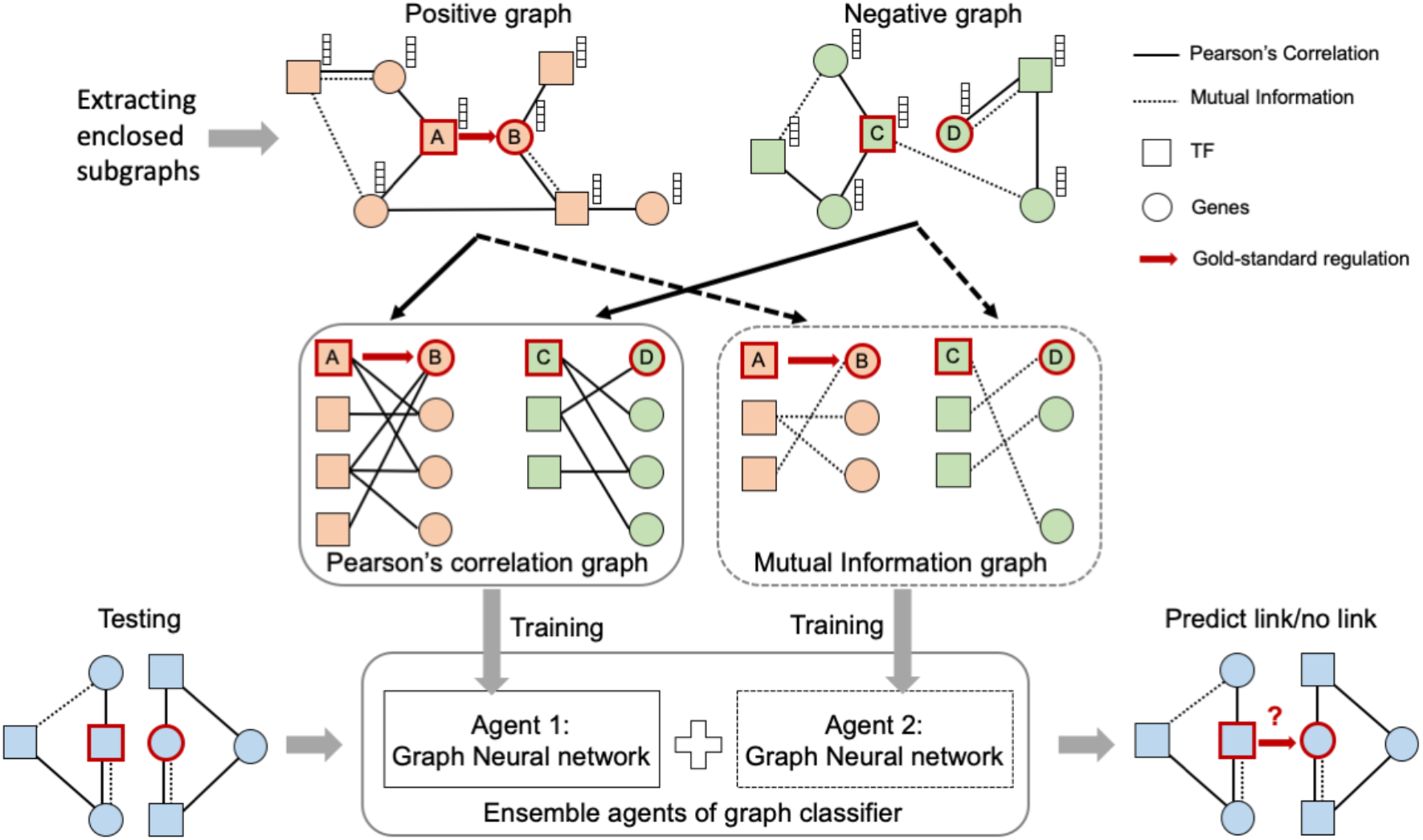
GRGNN scheme. Noisy starting skeletons derived from Pearson’s correlation and mutual information are used to generate the enclosed positive subgraph centering with A and B, and the negative graph centering with C and D. Graph neural networks as the agents are learned independently. An ensemble classifier is built upon these agents and used for the link prediction through graph classification.

### 2.1. Construct noisy starting skeletons

In order to incorporate the local structure of the input, heuristic methods are applied to infer relationships between TFs and their target genes through the input of gene expression *EX* in both training and testing datasets. Due to the limitation of existing heuristic methods, the inferred links are noisy, but integrating these links together as a starting skeleton can provide guidance in training.

In contrast to inherently unknown *GRN*, we define *GRN* ′ as the noisy skeleton inferred from the gene expression data. Totally *k* noisy skeletons *GRN*_*i*_ ′ =< *T,G, E*_*i*_ ′ >,*i* ∈[1, *k*]are constructed from *k* heuristic functions. Given TF *t* and gene *g*, each heuristic function *H*_*i*_ (*t, g*) ∈[0,1],*i* ∈[1, *k*]. The adjacent matrix in the *i-*th noisy skeleton is defined as:

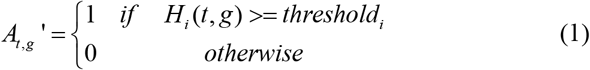

The thresholds are set as the parameters for tuning.

### 2.2. Extract enclosed subgraphs

Most of TF and target pairs are actually unlabeled with unknown regulatory information, and hence we predict them using co-expression at the first-order approximation for the graph topology *GRN*_*i*_ ′. For each of the known regulatory partners *t* ∈ *T* and *g* ∈{*T,G*}, (*t, g*) ∈ *E*, extract a subgraph *SG*_*i*_ (*t, g*)^+^ containing themselves and their *h*-hop neighbors on this noisy skeleton *GRN*_*i*_ ′ as the positive subgraphs. Meanwhile, randomly select *t* ∈ *T* and *g* ∈{*T,G*}, (*t, g*) ∉ *E*, extract a subgraph *SG*_*i*_ (*t, g*)^−^ containing themselves and their *h*-hop neighbors on the noisy skeleton *GRN*_*i*_ ′ as the negative subgraphs. Although such a negative set may contain false negatives due to undiscovered regulatory relationships, this is a widely used process of choosing negative examples. To get a balanced dataset, usually the number of negative links is chosen as the same size as the positive links.

### 2.3. Add node labels and features

A simple labeling function *label*(*i*) is used for marking node *i’* s roles in *SG*(*t, g*):

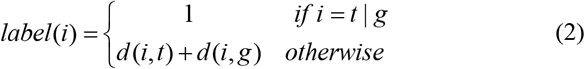

Only the centered target nodes *t* and *g* are labeled with 1, while the importance of nodes decreases when the node is far away from the center nodes. With appropriate labels, GNN can learn the structural information whether a link exists between the target nodes.

As either a TF or a target gene, each node in GRN has abundant information to reveal its biological roles. Generally, these features can be categorized into *explicit features* and *structural embeddings*. Only gene expression data are used to build node features. For gene expression vector *EX*_*i*_ of gene *i, i* ∈{*T,G*}, *μ* is the mean and *σ* is the standard deviation. *Q*_1_, *Q*_2_ and *Q*_3_ are quantiles of expression values. expression values. *Q*_0_ and *Q*_4_ are set as the minimum and maximum

After several experiments, gene expression features z-score∈(−∞, +∞) as Eq.3, *σ* and four Quantile Percentage ∈ (0,1) as Eq.4 along with TF∈{0,1} are defined as the explicit features.

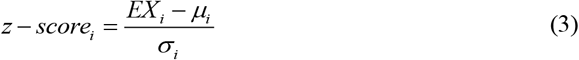

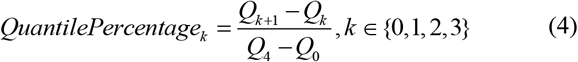

Graph embedding is a learned continuous feature representation for nodes in networks. *Node2vec* is applied to learn a mapping of nodes to a low-dimensional space of features that maximizes the likelihood of preserving network neighborhoods of nodes (Grover and Leskovec, 2016). Complementary to node labeling, graph embedding is aimed to capture the topological structure of the networks with the diversity of connectivity patterns in networks. The graph embedding is concatenated with explicit features together as the node feature vectors. For different *GRN*_*i*_ ′, the explicit features are consistently similar for all the nodes sharing the same gene expression input. These topological differences result in diverse node labels and graph embedding.

### 2.4. Build ensemble GNN classifiers

With the whole graph and the node features as the input, any GNN for graph classification could be used as a classifier. Here, DGCNN (Zhang *et al*., 2018) is used to address the graph classification, which adopts a quasi Weiseiler-Lehman subtree model (Shervashidze *et al*., 2011) to extract nodes’ local substructure features, and pool these nodes in order. Finally, a convolutional network work (CNN) is followed to read sorted graph representations and make predictions.

For each *GRN*_*i*_ ′ = (*A*_*i*_ ′, *EX*), where the adjacent matrix *A*_*i*_ ′ is built from gene expression *EX, k* GNN classifiers are built upon *k* sets of positive and negative enclosed subgraphs based on *k* heuristic functions. Then an ensemble classifier is built upon these *k* classifiers. Define *L* as the logits of the last layer of GNN with a softmax function, where *w1* and *w0* are neural weights for binary prediction:

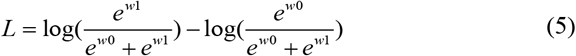

For *i* ∈[1, *k*], *α*_*i*_ is the weight, then the logits of the ensemble classifier *L*_*ensemble*_ can be defined as:

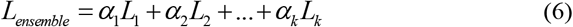

subject to *L*_1_ + *L*_2_ + … + *L*_*k*_ = 1 and *L*_*i*_ > 0,*i* ∈[1, *k*]. The parameter *α* can be trained either through a neural network or a simple least square regression.

### 2.5. Semi-supervised learning

A semi-supervised learning strategy is introduced to select a reliable negative sample set from the unlabeled datasets. Inspired by classical text classification S-EM (Liu *et al*., 2003), the basic idea is to build and maintain a Reliable Negative sample set RN through training iteratively. The process starts from randomly selecting samples from unlabeled data, the initial negative samples trained and tested by themselves are the initial RN. Keep RN and replace others with other unlabeled samples, and then train and test themselves iteratively. Each time keep negative samples as RN till equilibrium. It’s an Expectation-Maximization (EM) process and shown to be successful in many other classification applications.

### 2.6. GRGNN implementation

GRGNN is a versatile framework that fits for many alternatives in each step. In its implementation, two classical context relatedness measurements, Pearson’s correlation coefficient and mutual information are used to calculate links as a noisy skeleton to guide the prediction on the feature vectors of gene expression. In this setting, simply set *α*_1_ = 0.5 and *α*_2_ = 0.5 in the ensemble step already obtained good results. GRGNN is implemented with Pytorch (Paszke *et al*., 2017) and tested under Linux Ubuntu 16.04. The code for GRGNN is available at https://github.com/juexinwang/GRGNN.

## 3. Experimental Results

### 3.1 Dataset

In this study, three datasets from *In silico, E. coli* and *S. cerevisiae* in the DREAM5 challenge (Marbach *et al*., 2012) were used as the benchmark for evaluating GRGNN. The details of the DREAM5 datasets and the gold standard network of TF-target interactions are described in Table 1. From Table 1, *In Silico* dataset is quite different from *E. coli* and *S. cerevisiae* datasets in the scale of nodes, edges, average degree per TF, and average degree per node. In this paper, we only focus on the GRN inference performance on the *E. coli* and *S. cerevisiae* datasets.

**Table 1.** Details of DREAM5 datasets. Only *E. coli* and *S. cerevisiae* are used for the analysis.

| Species | # nodes | #TF | #Target Genes | # Links | # Samples | #avg degree per TF | #avg degree per node |
| --- | --- | --- | --- | --- | --- | --- | --- |
| *In Silico* | 1643 | 195 | 1448 | 4012 | 805 | 2.442 | 20.57 |
| *E. coli* | 4511 | 334 | 4177 | 2066 | 805 | 0.458 | 6.19 |
| *S. cerevisiae* | 5950 | 333 | 5617 | 3940 | 536 | 0.662 | 11.83 |

### 3.2 Comparing with supervised methods

We first compared GRGNN with other supervised methods in inductive performances. The model trained from *E. coli* was applied to predict regulatory relationships in *S. cerevisiae*, and the model trained from *S. cerevisiae* was used to predict *E. coli*. SVM and Random Forest (RF) are set as the baseline methods. GRGNN in *0*-hop is evaluated to get a fair comparison with the baselines, which means no neighbor information from the graph data structure is used with *0*-hop, other than the graph embedding. To quantify whether neighbors in the graph bring additional predictive power, *1*-hop GRGNN is evaluated along with *0*-hop GRGNN. GRGNN guided by both Pearson’s correlation coefficient and mutual information as the noisy starting skeleton is evaluated individually with their ensemble form GRGNN-EN. The cutoff of Pearson’s correlation coefficient is 0.8 and only mutual information larger than 3 *σ* is chosen as the guiding edge. For each of the evaluations, node features with only explicit features are compared with explicit features plus graph embedding learned from *node2vec*. As the input graphs are relatively small in scale, the dimension of the embedding feature vector here is set as 1. SVM and RF are implemented through python package *sklearn* (Pedregosa *et al*., 2011). Measurements such as *accuracy, precision, recall*, and Matthews correlation coefficient (*MCC*) are used to evaluate the performances. To evaluate performances of the proposed methods, negative samples are randomly selected as the gold-standard positive samples in both training and testing processes. All the experiments are run 5 times and take the mean and standard deviation.

Table 2 is the evaluation results on these balanced datasets both in *E. coli* and *S. cerevisiae*, which shows that ensembled GRGNN outperforms SVM/RF in GRN inferences in nearly all the criteria. Even though both GRGNN agents guided by noisy starting skeletons basically beat baselines in most cases, the ensemble of these two agents of GRGNN_PC and GRGNN_MI could persistently improve the results and help provide much more robust results. Furthermore, *1*-hop GRGNN outperforms *0*-hop GRGNN persistently in most of the criteria of both datasets, which indicates integrating neighbors brings more predictive power to the graph model. Even considering trained on two endpoints without neighbors as degraded with 0-hop, GRGNN outperforms SVM/RF with/without embedding features. This is due to the pooling procedure of GNN, where GNN itself outperforms SVM/RF. Even bringing some variances, adding graph embedding of the enclosed subgraph generally improves the performances for GRGNN. Especially, artificially involving graph embedding from the enclosed graph significantly improves the performances of the baseline SVM/RF, shows the power of neighbors. It could be explained as structural information from the noisy skeletons is involved as graph embedding in the training processes. In summary, GRGNN outperforms the baseline as it obtains predictive power from neighbor information through the guidance of noisy skeleton of embedding and ensemble processes.

**Table 2.** Evaluating inductive performance with supervised GRN inferring methods on balanced datasets. Feature E is the explicit expression features and G is graph embedding.

| Methods | Features | *E. coli* |  |  |  | *S. cerevisiae* |  |  |  | Note |
| --- | --- | --- | --- | --- | --- | --- | --- | --- | --- | --- |
|  |  | Accuracy | Precision | Recall | MCC | Accuracy | Precision | Recall | MCC |  |
| SVM | E | 0.621±0.000 | 0.628±0.000 | 0.594±0.000 | 0.242±0.000 | 0.505±0.000 | 0.557±0.000 | 0.056±0.000 | 0.026±0.000 | Baseline |
|  | G+E | 0.704±0.027 | 0.761±0.009 | 0.596±0.092 | 0.420±0.045 | 0.643±0.000 | **0.941±0.001** | 0.304±0.000 | 0.387±0.001 | Enclosed Graph + SVM |
| RF | E | 0.568±0.000 | 0.595±0.000 | 0.423±0.000 | 0.141±0.000 | 0.507±0.000 | 0.520±0.000 | 0.186±0.000 | 0.019±0.000 | Baseline |
|  | G+E | 0.635±0.031 | 0.807±0.057 | 0.359±0.070 | 0.326±0.061 | 0.658±0.004 | 0.848±0.012 | 0.384±0.007 | 0.377±0.009 | Enclosed Graph + RF |
| GRGNN_PC<br>(hop0) | E | 0.653±0.001 | 0.652±0.001 | 0.726±0.153 | 0.306±0.001 | 0.537±0.000 | 0.674±0.001 | 0.145±0.000 | 0.121±0.001 | - |
|  | G+E | 0.670±0.150 | 0.677±0.160 | 0.776±0.134 | 0.352±0.286 | 0.630±0.072 | 0.777±0.155 | 0.492±0.290 | 0.306±0.171 | - |
| GRGNN_PC<br>(hop1) | E | 0.586±0.007 | 0.580±0.007 | 0.625±0.009 | 0.173±0.014 | 0.566±0.000 | 0.662±0.002 | 0.395±0.280 | 0.164±0.000 | - |
|  | G+E | 0.696±0.078 | 0.677±0.062 | 0.773±0.100 | 0.395±0.160 | 0.655±0.059 | 0.746±0.121 | 0.518±0.078 | 0.343±0.115 | - |
| GRGNN_MI<br>(hop0) | E | 0.614±0.002 | 0.581±0.001 | 0.810±0.025 | 0.251±0.003 | 0.536±0.000 | 0.678±0.000 | 0.136±0.001 | 0.119±0.000 | - |
|  | G+E | 0.820±0.008 | **0.874±0.015** | 0.741±0.034 | 0.647±0.011 | 0.632±0.175 | 0.866±0.269 | 0.396±0.070 | 0.321±0.424 | - |
| GRGNN_MI<br>(hop1) | E | 0.652±0.003 | 0.635±0.002 | 0.718±0.019 | 0.306±0.006 | 0.534±0.001 | 0.571±0.001 | 0.326±0.117 | 0.079±0.002 | - |
|  | G+E | 0.767±0.068 | 0.744±0.077 | 0.847±0.025 | 0.540±0.134 | 0.566±0.202 | 0.695±0.283 | 0.579±0.149 | 0.150±0.453 | - |
| GRGNN_EN<br>(hop0) | E | 0.643±0.000 | 0.619±0.001 | 0.743±0.002 | 0.293±0.002 | 0.537±0.000 | 0.676±0.001 | 0.141±0.000 | 0.120±0.000 | - |
|  | G+E | 0.771±0.100 | 0.766±0.141 | **0.862±0.076** | **0.568±0.187** | 0.662±0.090 | 0.818±0.221 | 0.568±0.229 | 0.388±0.195 | Baseline Compared |
| GRGNN_EN<br>(hop1) | E | 0.656±0.000 | 0.637±0.000 | 0.730±0.000 | 0.318±0.003 | 0.570±0.000 | 0.630±0.002 | 0.340±0.002 | 0.158±0.001 | - |
|  | G+E | **0.809±0.033** | 0.743±0.069 | 0.853±0.112 | 0.564±0.153 | **0.684±0.056** | 0.770±0.147 | **0.574±0.083** | **0.393±0.135** | Proposed Method |

### 3.3 Comparing with unsupervised methods

The inductive capability of supervised methods makes it comparable with unsupervised GRN inferring methods. For GRN, supervise learning on the extremely unbalanced dataset brings strong bias in favoring negative samples. Take *E. coli* for example, there are 334×(4511−1) = 1,506,340 possible links in total, and only 2, 066 among them are confirmed gold standard positive links. Hence, a receiver operating characteristic (ROC) curve and precision-recall curve for all the methods on both *E. coli* and *S. cerevisiae* datasets were generated in Figure 2. We chose the widely accepted random forest based unsupervised method *GENIE3* as the representative unsupervised methods in comparison. GENIE3 outputs predicted links between TFs and target genes ranked by statistical confidences. In this study, the python implementation of GENIE3 is downloaded from its official GitHub repository. The default parameters were applied on both *E. coli* and *S. cerevisiae* datasets, and the top 1,000,000 predicted links were used for evaluation. To fairly compare unsupervised methods with supervised methods, GRGNN was learned purely from the *S. cerevisiae* dataset in the study of *E. coli*, and both GRGNN and GENIE3 were fed with gene expression data only from *E. coli* in testing (i.e., without using any TF-target gene labels for training). The same protocol proceeded in the study of *S. cerevisiae* with trained GRGNN from *E. coli*. All supervised methods were trained and tested on a balanced dataset. In this experiment, GRGNN used its ensemble version with *1*-hop neighbors and graph embedding. The baseline SVM used explicit features from genes only.

**Figure 2.**
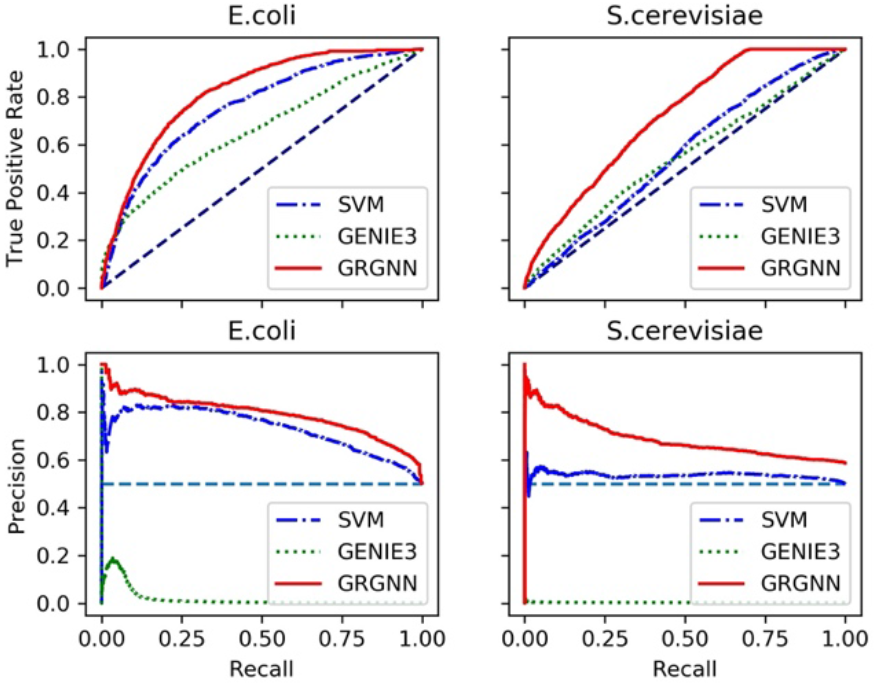
ROC curve and Precision-Recall curve on balanced training and testing

Figure 2 shows that GRGNN outperforms GENIE3 and baseline SVM methods on both ROC curve and Precision-Recall curve. Our results are consistent with existing works in supervised-unsupervised comparison in GRN, that supervised methods are typically superior (Maetschke *et al*., 2013). Besides, our results have demonstrated when training and testing in different datasets, GRGNN has better generalization capability inductively than GENIE3.

### 3.4 Inferring regulatory from a different number of layers

One common question in building the GNN models is how many layers of neighbors are sufficient for graph inference. An empirical test on dataset *S. cerevisiae* was processed by GRGNN. Starting from choosing no neighbors, *0*-hop GRN only relies on the pooling process on all node presentations to make the prediction. Then, layers and layers of neighbors were added into the models incrementally until reaching hop-9, which means in this case, the enclosing training and testing graph include far away nodes in distance up to 9 from the centered linked TF and target gene pairs.

Accuracy, precision, recall, and MCC are evaluated through these models in Figure 3, which indicates that the step adding *1*-hop to *0*-hop brings extra predictive power with the neighbors as the local structure in the graph. After that, adding more hops does not seem to bring significantly better results in GRN. This phenomenon may indicate that few hops of GNN contain almost all information for link prediction from its local structure in the graph, as the information of other parts of the network may be encoded well through graph embedding. In practice, *1*-hop GRGNN itself could get good results. Our results on GRN are consistent with the *γ* -decaying theory (Zhang and Chen, 2018), in which first-order and second-order heuristics can be perfectly computed from 2-hop enclosing subgraphs, while high-order global heuristics can be approximated from *h*-hop enclosing subgraphs with an exponentially smaller error.

**Figure 3.**
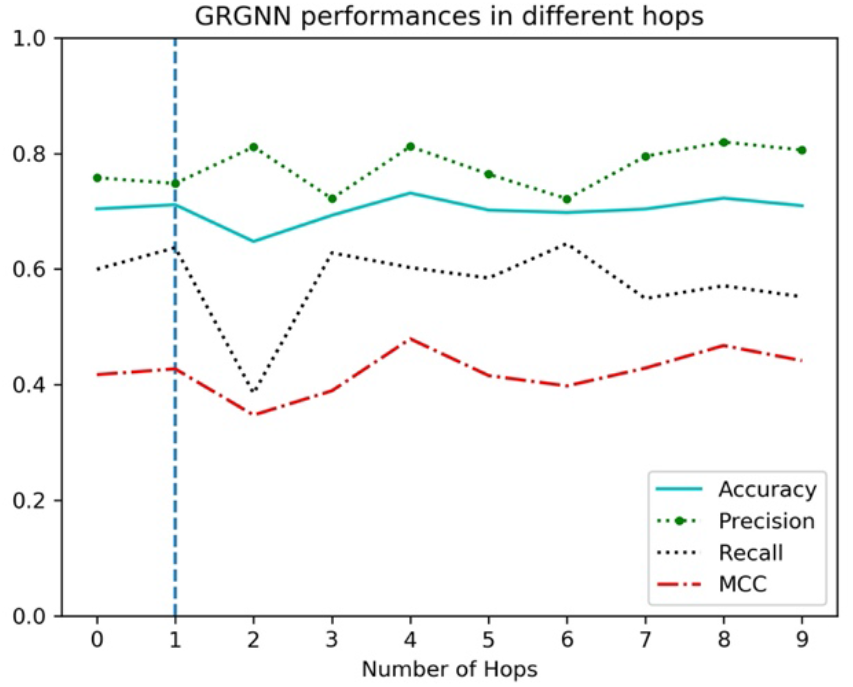
Performances of GRGNN in different numbers of hops.

### 3.5 Heuristic starting skeletons help regulatory inference

Considering the factor that links existing only in small proportion between the available nodes in our DREAM benchmark network, we generated random links between TFs and all the TF/targets in the uniform distribution with probability 0.003. This random network is used as the starting skeleton to replace the informative heuristic starting skeletons generated from Pearson’s correlation and mutual information. We run GRGNN 10 times with hop 1 in the same setting in Table 2. For *E. coli* without graph embedding features, the average and standard deviation of Accuracy, Precision, Recall, and MCC are 0.553±0.027, 0.570±0.044, 0.460±0.113, 0.112±0.058. For *E. coli* within graph embedding features, the average and standard deviation of Accuracy, Precision, Recall, and MCC are 0.571±0.039, 0.597±0.090, 0.574±0.206, 0.162±0.086. These weak prediction powers may only come from the endpoints, random networks as the starting skeleton brings random neighbors as the noises. Comparing with results in Table 2, we can see our usage of Pearson’s correlation and Mutual Information indeed brings useful information to the model.

## 4. Discussion

From the experiment’s results, the inductive prediction power of GRGNN on GRN may come from the following aspects. (1) **Ensemble of various heuristic skeletons**. Even a skeleton built from Pearson’s correlation coefficient or mutual information has a relatively low signal-to-noise ratio, an appropriate ensemble processes in the end alleviated these noises along with diverse information from different angles in linear correlation and information theory. Meanwhile, training and testing the same source of heuristics brings GNN opportunities to learn a mapping from the heuristic to the genuine regulatory relationships. (2) **Graph embedding captures network topological structures for link prediction**. Consistent with the biological hypothesis in GRN, subgraph with neighbors is much more informative than the regulatory pairs itself. Learned embeddings explored neighborhoods to have a better representation of the graph. This structural information may be used to explain why nearly every model obtained better performances when using graph embedding information. (3) **Carefully selected explicit features from gene expression**. Gene expression is the main input for GRN inferences. Comparing with learned embedding on noisy skeletons, gene expression data are the direct and dominant factors for relation inferences. To increase model generalization for different species and conditions, z-score, standard deviation, and quantile percentages are selected to describe the overall distribution and tendency of the input expression. (4) **GNN as the graph classifier**. Different from success in the fixed grid of image classification, a well-established convolution neural network cannot handle graph well. Advances in representation, convolution and pooling on the graph data structure in GNN make high quality graph classifier feasible. (5) **Biological meaning in the graph formulation**. Subtracting a local graph as the regulatory unit is supported by the network motifs hypothesis in transcription networks (Alon, 2007). The same network motifs have already been observed to conserve across diverse organisms. The formulation as a graph classification inherently meets the biological meaning of GRN.

The main limitation of this work is the datasets used. *E. coli* and *S. cerevisiae* are relatively well-studied small model species. These data are the only benchmark having systematically clear, experimental validated, gold standard regulatory relationships. Inductive goals may be easy to obtain on these two species. With the expansion of regulatory relationship identification and deeper understanding of the regulatory mechanisms, GRGNN can be trained and tested on more species such as human, mouse and plants. Furthermore, GRGNN is flexible for adopting different technologies in setting up a heuristic skeleton, incorporating structural features, and choosing different graph classifiers. For different purposes, it has great potential to test combinations of other embeddings with other cutting-edge classifiers such as DiffPool (Ying *et al*., 2018) and K-GNN (Morris *et al*., 2019).

## Acknowledgments

This work was supported by the National Science Foundation Plant Genome Program [#IOS-1734145 and #IOS-1546873], and the National Institutes of Health [R35-GM126985].

